# Finding the right tool: a comprehensive evaluation of microglial inducible cre mouse models

**DOI:** 10.1101/2023.04.17.536878

**Authors:** Alicia Bedolla, Gabriel Mckinsey, Kierra Ware, Nicolas Santander, Thomas Arnold, Yu Luo

## Abstract

The recent proliferation of new *Cre* and *CreER* recombinase lines provides researchers with a diverse toolkit to study microglial gene function. To determine how best to apply these lines in studies of microglial gene function, a thorough and detailed comparison of their properties is needed. Here, we examined four different microglial *CreER* lines (*Cx3cr1^CreER(Litt)^*, *Cx3cr1^CreER(Jung)^*, *P2ry12^CreER^*, *Tmem119^CreER^*), focusing on (1) recombination specificity; (2) leakiness - degree of non-tamoxifen recombination in microglia and other cells; (3) efficiency of tamoxifen-induced recombination; (4) extra-neural recombination -the degree of recombination in cells outside the CNS, particularly myelo/monocyte lineages (5) off-target effects in the context of neonatal brain development. We identify important caveats and strengths for these lines which will provide broad significance for researchers interested in performing conditional gene deletion in microglia. We also provide data emphasizing the potential of these lines for injury models that result in the recruitment of splenic immune cells.

## Introduction

Microglia, the resident macrophages of the neural parenchyma, regulate a variety of processes necessary for brain development, homeostatic function, and injury/disease response. In addition to their role in innate immunity in the brain and surveillance of parenchyma, during both development and adulthood, microglia have been credited with synaptic pruning which plays a vital role in synaptic plasticity^1–4^. Outside the realms of development and homeostasis, microglia also respond to disease and injury, with varying activation profiles depending on the challenge^5,6^. With the increase in knowledge and recognition of these processes has come a significant expansion in the number of *Cre* and *CreER* recombinase mouse lines to enable precise genetic targeting of microglia^7–10^. Manipulations of the fractalkine receptor *Cx3cr1* gene locus have been one of the primary methods used to drive *Cre* and *CreER* expression in microglia (reviewed in^11^). Indeed, *Cx3cr1* is strongly expressed in microglia as well as other myeloid subsets, providing relatively specific gene targeting. Since their establishment, these lines have been collectively utilized in over 1000 published reports. These reports highlight the broad utility of *Cx3cr1*-based gene targeting but also describe several potentially important drawbacks including non-microglia recombination (lack of specificity)^12^, leakiness^13^, and off-target effects^12^. Based on then-emerging transcriptomic profiling, *Sall1^CreER^* mice^14^ were proposed to be more specific for the genetic manipulation of microglial cells than *Cx3cr1^CreER^* mice. However, a detailed characterization later found that *Sall1^CreER^* recombines neural-ectodermal lineages including neurons, astrocytes and oligodendrocyte populations, and also has significant tamoxifen-independent (leaky) recombination in microglia^12^. Furthermore, *Sall1^CreER^*, like *Cx3cr1^CreER^* mice, was generated as a knock-in/knock-out at the endogenous gene locus. As Sall1 is known to be critical for the maintenance of microglial homeostasis and activation^14,15^, heterozygous loss of *Sall1* in -CreER mice could have important impacts on microglia complicating interpretation of lineage tracing and gene knockout experiments.

In recent years, several new *CreER* mice were generated to overcome the drawbacks of existing microglia-targeting lines. In particular, *Tmem119^CreER^* and *P2ry12^CreER^* lines were made to have increased specificity for microglia while sparing brain border macrophages, such as perivascular and pial macrophages. In addition to this increased specificity, these new lines lack recombination in circulating monocytes, making it easier to distinguish microglia from invading monocytes in the context of injury or disease. With the advent of these new lines, a systematic and direct comparison of their properties is needed. A number of characteristics are of particular interest to researchers when deciding which *CreER* to use for conditional mutagenesis studies including: (1) specificity - what cells in CNS/brain are recombined by the *CreER*; (2) leakiness - the degree of non-tamoxifen recombination in microglia and other cells; (3) efficiency - how well loxP-flanked DNA regions are recombined in the presence of tamoxifen; (4) extra-neural recombination - related to specificity; the degree of recombination in cells outside the CNS, particularly myelo/monocyte lineages, that can enter the CNS in development or disease; (5) off target effects - whether there are unintended of gene targeting and/or tamoxifen administration in particular lines. With these five primary characteristics in mind, we performed a rigorous evaluation of four publicly-available (Jackson Laboratories) microglia-targeting *CreER* lines: *Cx3cr1^CreER(Litt)^* (generated from the Littman lab^16^), *Cx3cr1^CreER(Jung)^* (generated from the Jung lab^17^), *Tmem119^CreER^* ^10^, and *P2ry12^CreER^* ^8^. We report that there is significant variability across lines in their leakiness and in their ability to recombine floxed alleles which we show is related both to loxP distance and intrinsic CreER activity or expression. We describe our unsuccessful efforts to improve recombination efficiency by crossing to the new iSuReCre mouse line. We also document significant differences in splenic recombination in the studied CreER lines, reflective of potential injury-induced monocyte recruitment from the spleen. In total, these comparative analyses provide important data for research groups eager to determine whether a particular microglial recombining *CreER* line would be appropriate and useful for studies of cell lineage tracing and/or conditional gene mutation in microglia.

## Results

### Different degrees of leakiness in microglia CreER lines

We first looked to confirm microglia specificity while also assessing leakiness in *P2ry12^CreER^* and *Tmem119^CreER^* mice. We analyzed *Cx3cr1^CreER(Jung)^*and *Cx3cr1^CreER(Litt)^*, both previously well characterized, as baseline controls. For these studies, we took advantage of what is widely regarded as the most sensitive Cre-recombinase reporter, *Ai9/Ai14*, and a less sensitive Cre reporter, *ROSA26-YFP* (hereafter referred to as *R26-YFP*) for comparison. Note that because *Cx3cr1^CreER(Litt)^* line carries an internal ribosome entry site (IRES) - enhanced yellow fluorescent protein (EYFP) gene reporter, we only examined a cross with the *Ai9-tdTomato* reporter in this line as *Cx3cr1^CreER(Litt)^Ai9*. We assessed mice treated with tamoxifen (180mg/kg daily gavage for 5 days), compared to mice treated with vehicle (sunflower oil + ethanol) kept in separate cages throughout the induction and analysis period (to ensure no tamoxifen contamination). Vehicle treatment provides a control to tamoxifen treatment and also allows for an assessment of leakiness – the number and types of cells recombined in the absence of tamoxifen.

As expected, tamoxifen induction in adult *Cx3cr1^CreER^* (Jung and Littman) mice resulted in recombination of the *Ai9 tdTomato* Cre reporter allele in microglia as well as brain border macrophages in the choroid plexus, meninges, and perivascular spaces (Fig 1 bottom, Fig S1). In contrast, tamoxifen treated *Tmem119^CreER^* and *P2ry12^CreER^* mice had more specific tdTomato labeling of microglia and no apparent labeling of brain border macrophages (Fig S1) or other parenchymal neurons and glia (not shown). *Tmem119^CreER^* lacked recombination of all choroid plexus macrophages, including Kolmer’s epiplexus cells which are targeted by *P2ry12^CreER^* and both *Cx3cr1^CreER^* lines (Fig S1). Interestingly, *Tmem119^CreER^* recombined numerous Iba1-negative cells in the choroid plexus, meninges, and around large blood vessels, which likely represent brain border fibroblasts known to strongly express *Tmem119*. Comparing recombination in mice with the *Ai9* versus the *R26-YFP* recombination reporter, we observed significantly fewer recombined Iba1+ brain macrophages in *Tmem119^CreER^* and *P2ry12^CreER^* lines with *R26-YFP* (94.55% for the *Tmem119^CreER^Ai9* mice and 95.81% for the *P2ry12^CreER^Ai9* mice vs 33.78% for *Tmem119^CreER^R26-YFP* and 36.51% *for P2ry12^CreER^Ai9*) (Fig 1; note that YFP fluorescence was enhanced using anti-GFP immunostaining to minimize differences in endogenous florescence). This finding was confirmed using FACS analysis (Fig S2). We attribute this difference in Cre reporting to the intrinsic sensitivity of the two Cre recombinase reporters; a function of inter-loxP distances (0.9 kb in *Ai9*; 2.7 kb in *R26-YFP*), mRNA stabilizing elements (*Ai9* has WRPE), enhanced promoter elements (*Ai9* uses a strong CAG promoter), and native fluorescence intensity (*Ai9* tdT vs YFP).

**Figure 1.**
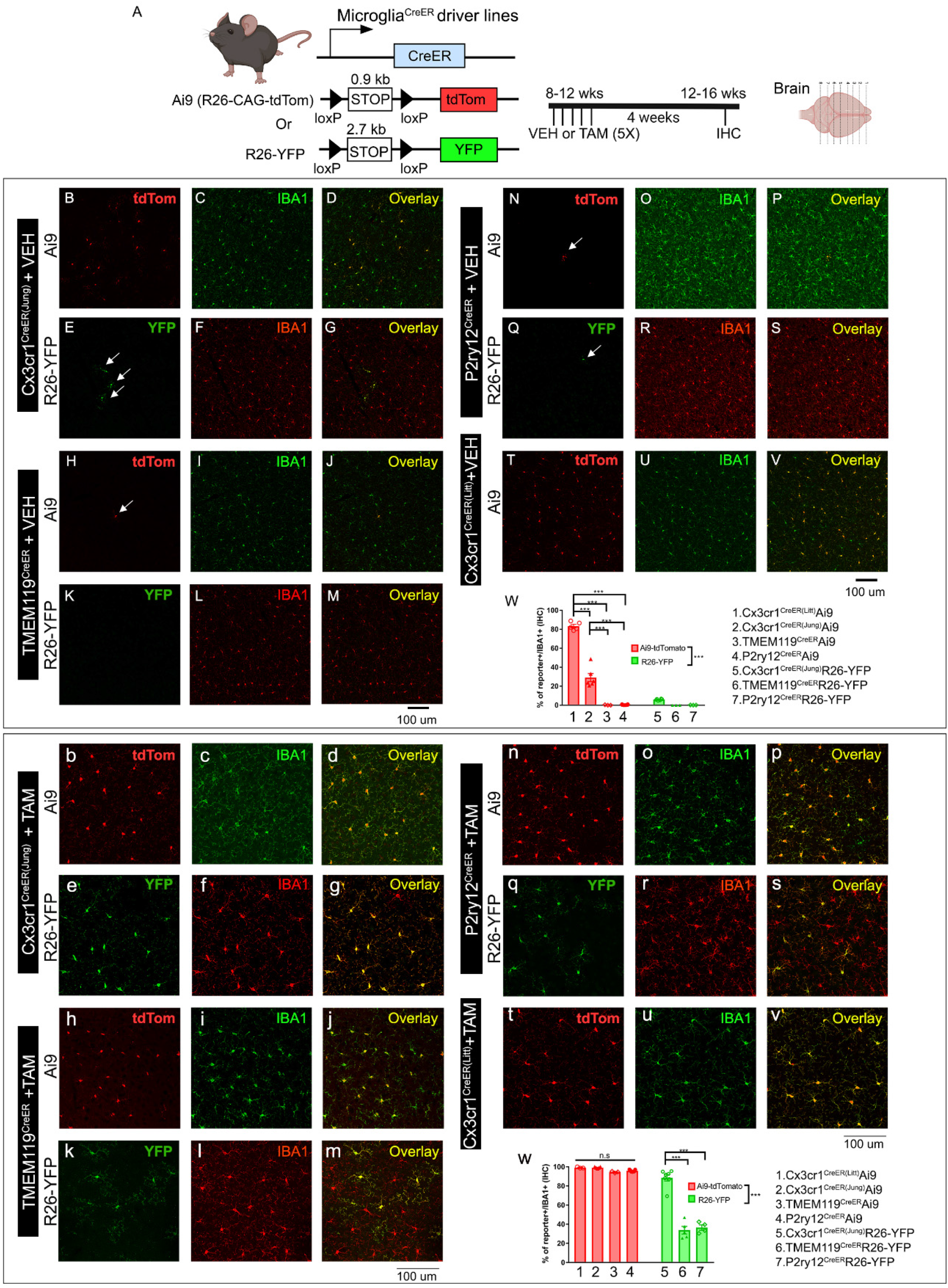
Evaluation of the TAM-independent leakiness and the efficiency of TAM-dependent cre recombination in the four different creER driver lines using either the Ai9 (tdTomato) or R26-YFP reporter mouse lines. The experimental timeline is shown in panel (A). Representative images from each cre driver and reporter line (B-V for VEH treatment and b-v for TAM treatment). Cre driver and the reporter line are indicated on the left side of the panels. Quantification of reporter+ cells in the IBA1+ populations in the brain is shown in panel W (for VEH treatment) and w (for TAM treatment). Representative images are taken from the cortical region which reflects the general and homogenous trend in the whole parenchyma. Each data point represents the average of 1 animal (the average for each animal is obtained by quantifying multiple brain sections at similar anatomical location) and the average for each animal was used as a single data point for statistical analysis. **p < 0.01 and ***p < 0.001, for Two way ANOVA analysis, Tukey post-hoc pair wise analysis. Ai9 vs R26-YFP is significantly different as a factor (p<0.001). Data were combined from 2 independent cohorts of mice. Scale bar: 100 μm. Compared to the two Cx3cr1^CreER^ lines (Littman and Jung), TMEM119^CreER^ and P2ry12^CreER^ show less leakiness in the absence of TAM but a decreased recombination efficiency and mosaic recombination in microglia.

Tamoxifen-independent tdTomato recombination was previously described in *Cx3cr1^CreER(litt)^* ^18^, *Cx3cre1^CreER(Jung)^* ^12^ and *P2ry12^CreER^* ^19^ (and unpublished observations) lines. Using the Ai9 Cre-reporter, we similarly observed relatively high degrees of tamoxifen-independent recombination in microglia in the two *Cx3cr1^CreER^* lines (82% and 28%, respectively), and sparse recombination (tdTomato expression) of microglia in *P2ry12^CreER^Ai9* and *Tmem119^CreER^ Ai9* mice in the absence of tamoxifen (Fig 1, top). Similar to the *P2ry12^CreER^Ai9* mice, we observed sparse non-tamoxifen recombination of microglia in *Tmem119^CreER^Ai9* mice. Using the *R26-YFP* reporter line, we observe much less tamoxifen-independent expression of the YFP reporter in the *Cx3cr1^CreER(Jung)^R26-YFP* brain and observed almost no YFP+ cells in *Tmem119^CreER^* or *P2RY12^CreER^R26-YFP* mice (occasionally one YFP^+^ cell in the whole brain section). Taken together, these data confirm the previously characterized microglia-specific recombination *Tmem119^CreER^* and *P2ry12^CreER^*, and document low rates of tamoxifen-independent Cre recombination (leakiness) in both of these lines as well as both *Cx3cr1^CreER^* lines, with the greatest relative leakiness observed in the *Cx3cr1^CreER(Litt)^* line.

### Gene targeting efficiency is related both to intrinsic CreER expression/activity and inter-loxP target length

Our data above show that the *Tmem119^CreER^* and *P2ry12^CreER^* lines are able to achieve highly efficient recombination in the *Ai9* allele (95% and 96%, respectively), whereas the recombination efficiency in the *R26-YFP* allele is comparatively less (34% and 37%, Fig 1 bottom). Similarly, the rates of tamoxifen-independent recombination in the *Cx3cr1^CreER(Jung)^* line is lower with the *Ai9* versus the *R26-YFP* allele (99% vs 89%). We considered whether this apparent difference in reporting is due to inter-loxP distance (or possibly other differences in reporter gene expression, stability, or fluorescence intensity) and not genomic location or DNA accessibility since both the *Ai9* and the *R26-YFP* reporter gene are inserted into the R26 loci. We therefore generated *P2ry12^CreER^Ai9/R26-YFP* double reporter mice and examined the number and relative intensity of recombined cells on a single cell basis using flow cytometry and immunohistochemistry (Fig 2). We observe that tdTomato+ cells are more abundant among reporter positive cells (% of tdTomato^+^ in all reporter^+^ cells = 82.52% versus YFP^+^ cells in all reporter^+^ cells= 41.55%. p<0.001, Student’s t-test, Fig 2 and Supplementary Table 1). However, there are also YFP^+^ cells that are negative for the tdTomato reporter (average = 17.48% of the total reporter+ cells, Fig 2). These results suggest that although the shorter length floxed *Ai9* reporter allele has a higher probability of recombination on a populational level, the recombination of the two different alleles in the same genomic loci is independently regulated; in any single cell, which allele is recombined is independent from the recombination of the alternate allele and does not always follow the size rule. Similarly, we speculate that recombination of a single copy of reporter allele does not always guarantee the recombination of both floxed alleles of any given target gene, especially when using a floxed reporter allele that has a short floxed region such as *Ai9* or *Ai14*.

**Figure 2.**
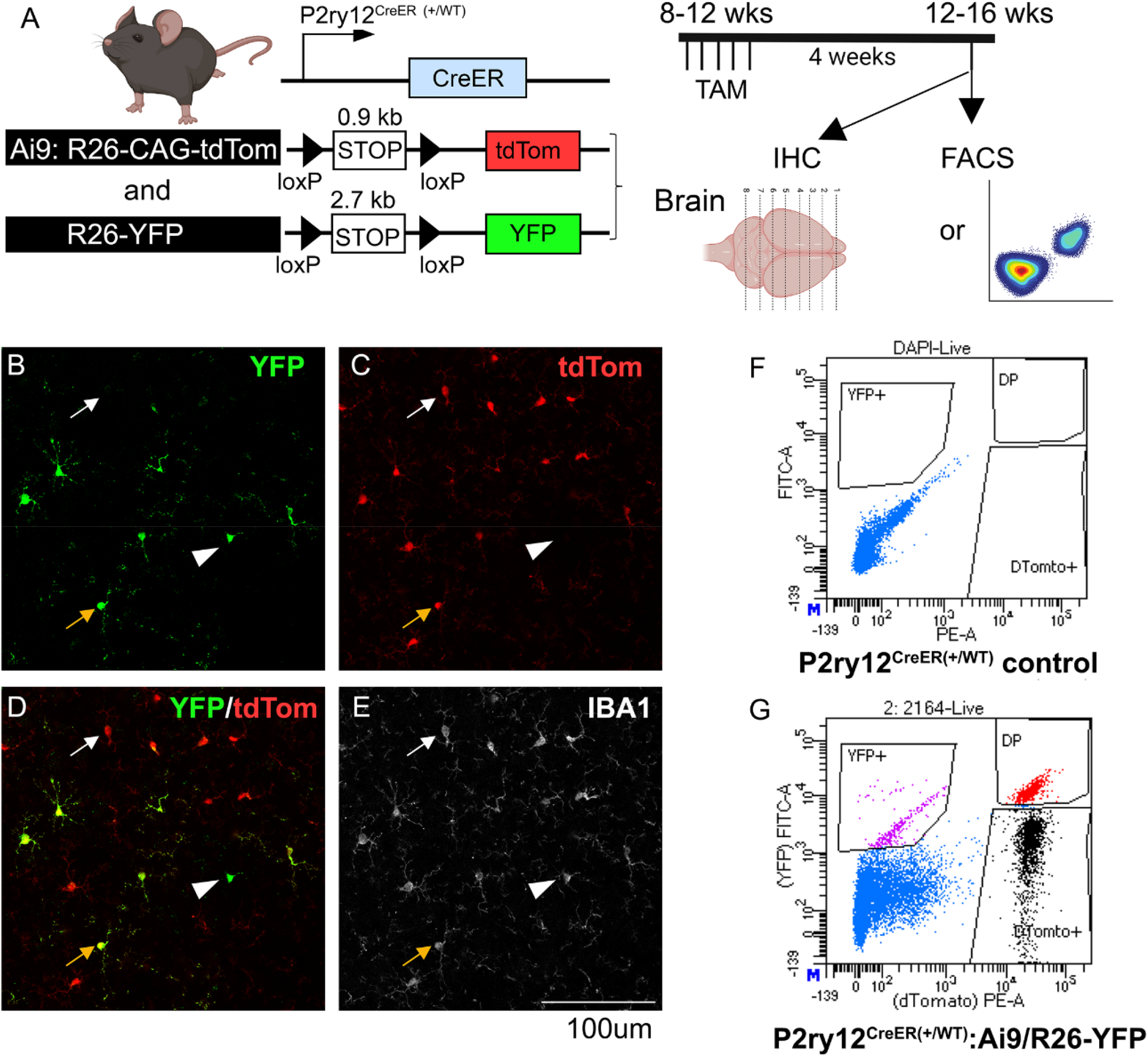
Independent recombination of the two floxed alleles on a single cell level in the P2RY12^CreER^ double reporter (Ai9:R26-YFP) mouse line. Evaluation of the TAM-dependent recombination of either Ai9-tdTomato or R26-YFP allele which are both located in the ROSA26 loci in a double reporter mouse in the P2RY12CreER line suggests that although on a populational level, Ai9-tdTomato reporter has a higher probability of being recombined than the R26-YFP allele, on a single cell level, the recombination of each individual allele can be independent and does not always follow the size of the floxed region rule. (A), experimental timeline. (B-E), IHC evaluation of the recombination of microglia on either of the reporter expression. Note tdTomato+:YFP+ double positive microglia (orange arrow), more abundant tdTomato+:YFP-microglia (white arrow) and the less abundant YFP+:tdTomato-microglia (white arrowhead). (F-G), representative FACS plots for no color control or the double reporter flow analysis. For quantification of % of each population, see Supplementary Table 1.

To more directly investigate the efficiency of microglial recombination, we generated different combinations of *Cx3Cr1^CreER(Jung)^* (apparent “strong” Cre) or *P2ry12^CreER^* (apparent “weak” Cre), bred to the following floxed alleles with varying inter-loxP distances (bred to homozygosity): *Tgfb1^fl/fl(ex3)^* (< 0.5kb loxp distance), *Alk5^fl/fl(ex3)^* (1.6-1.7 kb loxP distance). Mice were bred with a recombination reporter (*R26-YFP* or *Ai9/Ai14*) to facilitate isolation of microglia cells that have undergone at least one recombination event. Total RNA was extracted from sorted microglia cells in all mouse lines, followed by cDNA library preparation and quantitative realtime PCR using allele-specific forward and reverse primers spanning the floxed and neighboring exon. Absence of gene amplification therefore indicates deletion of floxed exons.

Our results (Fig 3) show that the *Cx3Cr1^CreER(Jung)^* line leads to significant and almost complete loss of the floxed exon in total RNA content for both *Cx3cr1^CreER(Jung)^Tgfb1^fl/fl^*and *Cx3cr1^CreER(Jung)^Alk5^fl/fl^* mice (Fig 3, 0.0024% for *Tgfb1* mRNA level compared to control mice, and 3.9% for *Alk5* mRNA level). *Iba1* mRNA levels in the sorted YFP+ microglia are not altered from the *Cx3cr1^CreER(Jung)^Tgfb1^fl/fl^* mice nor the *Cx3cr1^CreER(Jung)^Alk5^fl/fl^* lines, indicating similar purity of sorted microglia and that gene deletion is specific to the targeted floxed allele. As *Cx3Cr1^CreER(Jung)^* mice recombine the *R26-YFP* reporter allele in most microglia (>88%), this result reflects a highly efficient deletion of the target exons. In contrast, *P2Ry12^CreER^*-mediated recombination resulted in a ∼50% reduction in both the *Tgfb1* and *Alk5* floxed gene regions. Because *P2ry12^creER^* recombines ∼ 30% of the total microglia cells at the *R26-YFP* locus, and our sorting strategy enriches for YFP^+^ cells, this efficiency rate is likely overestimating true recombination efficiency at the gene of interest. Indeed, in *P2ry12^CreER^Alk5^fl/fl^-Ai9* (with the easier to recombine *Ai9* reporter), >95 % of sorted IBA1^+^ microglia were tdT^+^, while *Alk5* mRNA is 84% of WT (Fig 3), indicating a lower overall recombination efficiency in P2ry12^CreER^ mice compared to the *CX3cr1^CreER(Jung)^* line. Taken together, our data suggest that both intrinsic CreER activities as well as target inter-loxP distance are important determinants in gene excision. Our results also highlight critical differences in apparent gene targeting efficiency when using various reporters, specific assessment of allele-specific gene loss, or methods to isolate/analyze a conditional gene targeting experiment (e.g. FACS versus immuno-histology).

**Figure 3.**
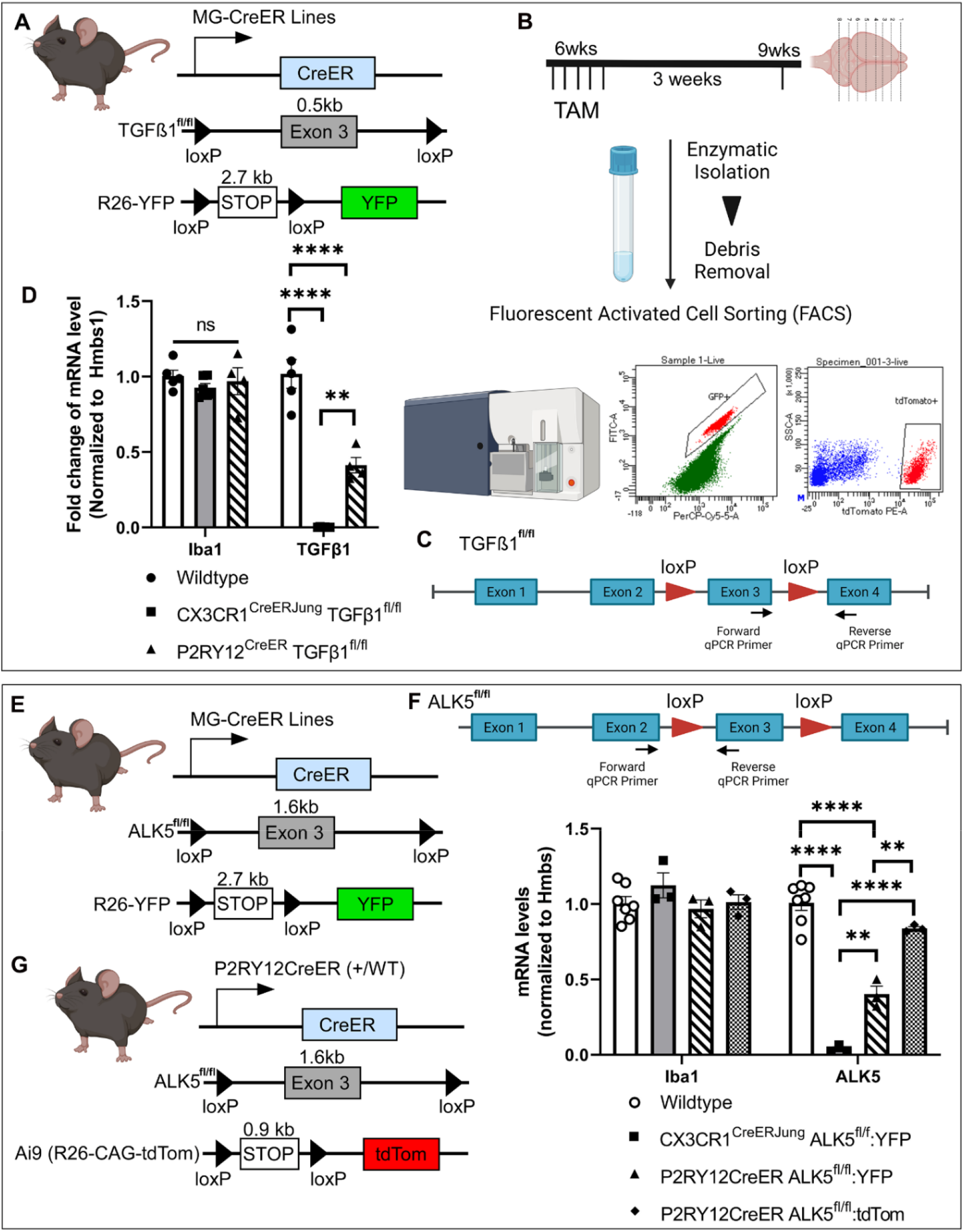
Evaluation of the gene deletion efficiency on distinct homozygous floxed target gene alleles in the Cx3cr1CreER(Jung) and P2Ry12CreER drivers using quantitative RT-PCR. (A-D). Animal genotype and experimental flow. Total mRNA levels are evaluated for the floxed exon in the TGFb1 gene in YFP+ microglia sorted from the Cx3cr1^CreER(Jung)^ or P2RY12^CreER^—TGFb1 ^fl/fl^-R26-YFP mice at 3 weeks after TAM treatment. (E-G). Total mRNA levels are evaluated for the floxed exon in the ALK5 gene in either YFP+ microglia sorted from the Cx3cr1^CreER(Jung)^ or P2RY12^CreER^—ALK5 ^fl/fl^-R26-YFP mice or tdTomato+ microglia from the P2RY12^CreER^—ALK5 ^fl/fl^-Ai9 mice at 3 weeks after TAM treatment. Each data point represents the average of 1 animal (the average for each animal is obtained by averaging 3 technical replication of the qRT-PCR reaction for that animal) and the average for each animal was used as a single data point for statistical analysis. **p < 0.01 and ***p < 0.001, **** p<0.0001 for Two way ANOVA analysis, Tukey post-hoc pairwise analysis.

### Lack of iSuReCre recombination by *P2ry12CreER*

Given the relatively weak recombination efficiency in *P2ry12^CreER^*, we were interested in whether we might enhance recombination while maintaining cellular specificity. The new iSuReCre mouse line, with Cre-inducible expression of both Cre-recombinase and membranous (Mb) Tomato reporter (Fig 4a), was made to accomplish this exact goal^20^. We generated *P2ry12^CreER^*iSuReCre mice, treated them with TAM at 4-8 weeks of age, then harvested and analyzed brains 2-4 weeks later. Our results (Fig 4) show that addition of the iSuReCre allele failed to label any microglia with the MbTomato reporter in the *P2ry12^CreER^* line. Interestingly, there was sparse MbTomato labeling of neurons in the cortex and striatum of *P2ry12^CreER^*iSuReCre mice which did not receive tamoxifen (Fig 4b-i), likely representing ectopic TAM-independent and cre-independent expression of MbTomato from the iSuReCre locus since we never observed neuronal recombination in *P2ry12^CreER^R26-YFP* or *P2ry12^CreER^Ai9* mice. To examine whether the iSuRe mouse model works properly in other non-microglia brain-specific Cre drivers, we generated a DCX^CreER^iSuReCre mice as a positive control (Fig 4j). *DCX^CreER^* mice target immature DCX^+^ neuroblasts in the dentate gyrus which later develop into mature NeuN^+^ neurons 4 weeks after administration of TAM. Indeed, brains from *DCX^creER^*iSuReCre mice had strong MbTomato expression in DCX^+^ neuroblasts and mature NeuN^+^/DCX^(-)^ neurons at 5days (Fig 4k-n) and 4 weeks (Fig 4o-r) after TAM administration, respectively. These data confirm that the iSuReCre allele is recombined by DCX^CreER^, similar to other Cre and CreER lines tested in the original publication^20^, but is not recombined in adult microglia by *P2ry12^CreER^*.

**Figure 4.**
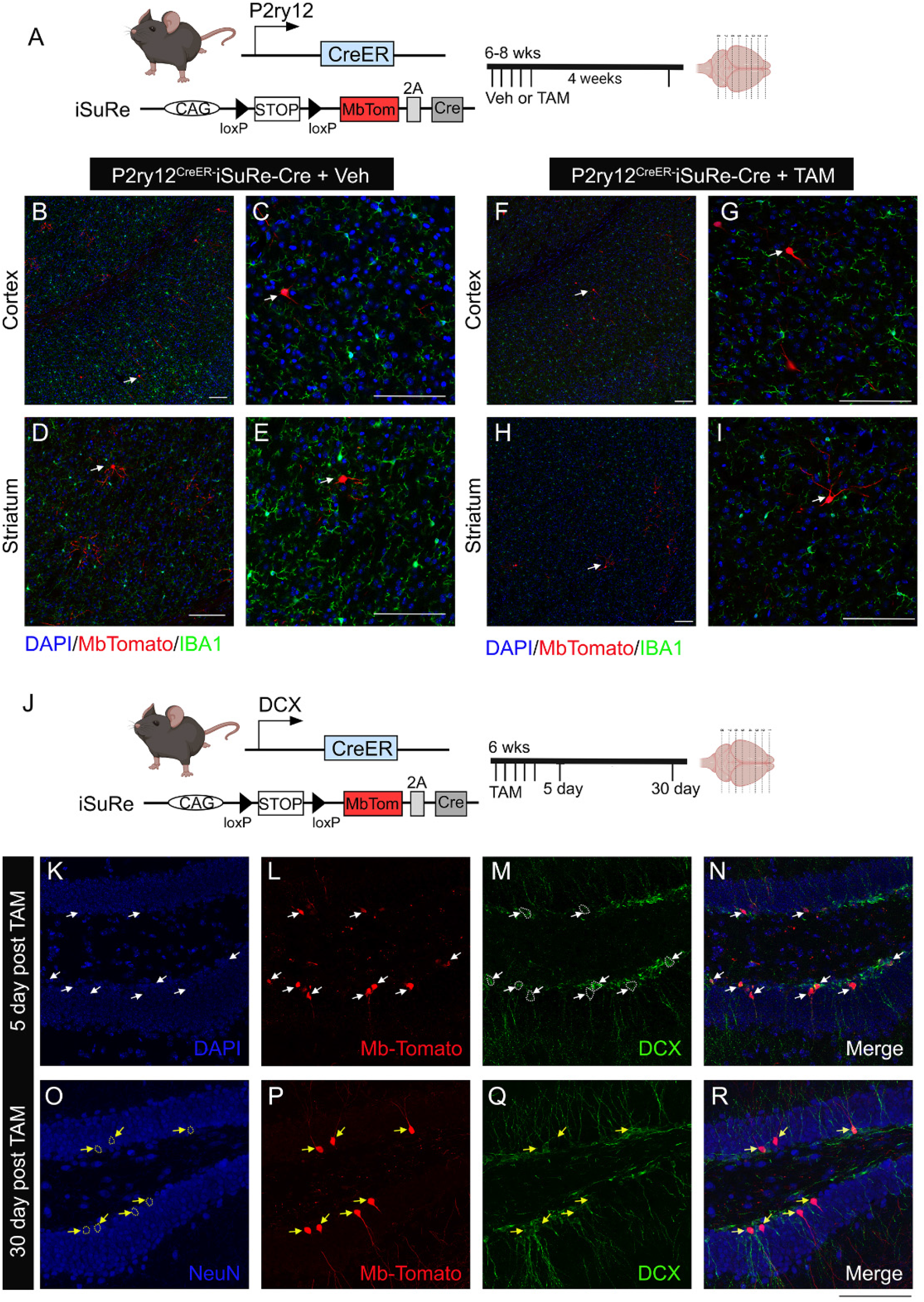
iSuRe-Cre mouse line successfully induces constitutive cre-P2A-MbTomato expression in DCX^CreER^ mouse line but not in the P2ry12^CreER^ mouse line after TAM treatment. (A) Illustration of mouse transgene constructs and experimental timeline. (B-E) in the absence of TAM, there is no MbTomato expression in microglia with ectopic MbTomato expression in cells that demonstrates typical neuron morphology in cortex and striatum. (F-G), treatment of TAM in mice does not induce MbTomato expression in microglia and presents with similar neuronal ectopic expression of MbTomato. (J-R). In contrast, in the DCXC^reER^-iSuReCre mice that are treated with TAM, at 5 days post TAM treatment, DCX+ immature neuroblasts are labeled with MbTomato protein (white arrows) and at 30 days post TAM treatment, MbTomato expression are mostly detected in DCX-NeuN+ mature neurons (yellow arrows), supporting that the iSuRe-Cre construct is able to be induced in a cohort of immature neuroblasts which mature later into NeuN+ neurons in the dentate gyrus of adult mice. Scale bar=100 µm.

### Persistent recombination of splenic macrophages in microglia CreER lines

One potential downside to using *Cx3cr1^CreER^*-based strategies in lineage tracking and conditional mutagenesis experiments is these lines’ known recombination of circulating blood cells, and the potential for these cells to invade and take residence in the CNS. With this in mind, strategies were developed to specifically label brain microglia vs peripheral monocytes/macrophage followed by a waiting period of > 3 weeks after TAM administration to allow for the turnover of circulating cells by non-recombined Cx3cr1-negative bone marrow progenitors^21,22^. However, these strategies are cumbersome or not feasible in developmental and injury contexts in which recombination and engraftment occurs more rapidly than the 3 week “washout period” and might include cells derived from non-myeloid (non-marrow) sources. The spleen is a particularly important site for the storage and rapid deployment of monocytes in the setting of development and inflammation^23–25^ and can be used as a surrogate for recombination circulating monocytes.

To this end, we examined the expression of recombination reporter genes in the spleens (matched to brain recombination, Fig 1) 4 weeks after TAM or vehicle treatment (Fig 5). In *Cx3cr1^CreER(Litt)^Ai9* mice, we observed substantial tamoxifen-independent base level tdTomato and YFP reporter expression in IBA1^+^ splenic macrophages, and a strong increase in reporter recombination/expression with persistence of these cells 4 weeks after tamoxifen administration, mirroring tamoxifen-independent and -dependent reporter recombination in the brain (Fig 1). We observed recombination of splenic macrophages in *P2ry12^CreER^Ai9* mice comparable to *Cx3cr1^CreER^* lines (as was previously reported), but less recombination of splenic macrophages in *Tmem119^CreER^* mice (Fg5h-m). Interestingly, there was strong recombination of non-myeloid (IBA1-negative) cells which, by comparison to the brain, may represent splenic adventitial fibroblasts^26^. These data highlight a relatively high degree of recombination in non-microglial splenic macrophages in TAM-treated (and untreated) *Cx3cr1^CreER^*and *P2ry12^CreER^* mice, representing a previously un-explored reservoir of recombined cells that could replace microglia and complicate the interpretation of lineage tracing and gene deletion experiments using these lines.

**Figure 5.**
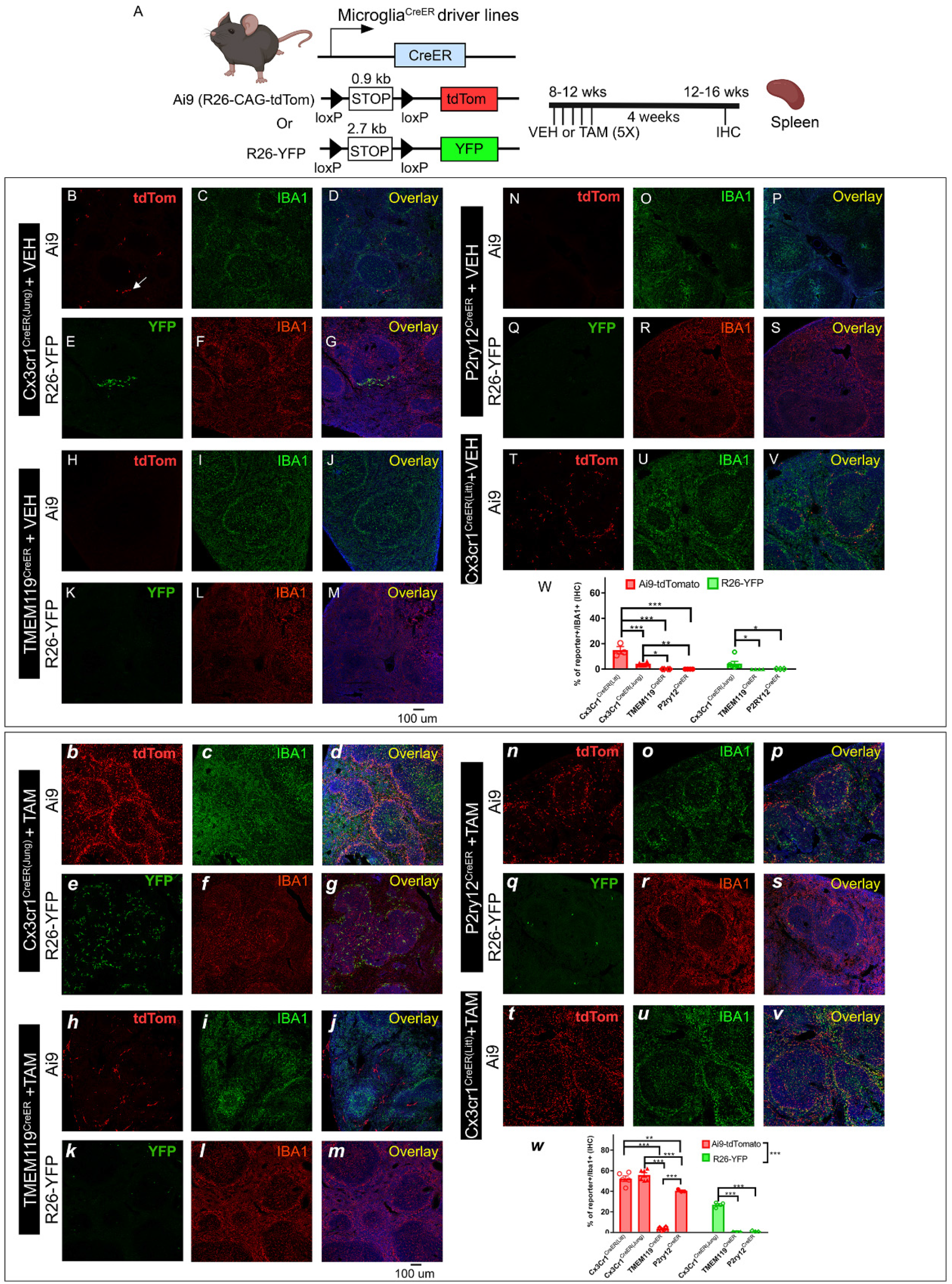
Evaluation of the splenic TAM-independent and TAM-dependent cre recombination in the four different creER driver lines using either the Ai9 (tdTomato) or R26-YFP reporter mouse lines. The experimental timeline is shown in panel (A). Representative images from each cre driver and reporter line (B-V for VEH treatment and b-v for TAM treatment). Cre driver and the reporter line are indicated on the left side of the panels. Quantification of reporter+ cells in the IBA1+ populations in the spleen is shown in panel W (for VEH treatment) and w (for TAM treatment). Each data point represents the average of 1 animal (the average for each animal is obtained by quantifying multiples pleen sections) and the average for each animal was used as a single data point for statistical analysis. **p < 0.01 and ***p < 0.001, for Two-way ANOVA analysis, Tukey post-hoc pairwise analysis. Ai9 vs R26-YFP is significantly different as a factor (p<0.001 for TAM treated group). Data were combined from 2-3 independent cohorts of mice. Scale bar: 100 μm.

### Off target effects of tamoxifen induced microglia-CreER expression

A recent evaluation of the *Cx3cr1^CreER(Litt)^* line found that neonatal tamoxifen exposure resulted in microglia with reduced expression of homeostatic genes, and reciprocal induction of activation genes related, in part, to interferon signaling^27^. Notably, *Tmem119^CreER^* did not show this same Cre-mediated non-specific effect in early postnatal microglia, and administration of tamoxifen to adult *Cx3cr1^CreER(Litt)^* mice showed no major phenotype. Given the high efficiency of *Cx3Cr1^CreER^*mouse lines in targeting microglia on a populational level and likely still popular usage in future microglia gene knockout studies, we looked to determine whether this same phenotype existed in other microglia-targeting CreER lines.

*Cx3cr1^CreER^*+ (Jung and Littman), and *P2ry12^CreER^*+ mice were intercrossed with wild type mice, and newborn pups were given tamoxifen (50ug dose IG or 500ug dose IP) on postnatal day P1,2,3 or P4,5,6 and then brains were harvested and analyzed on P15 or P30 (Fig 6). For convenience we used brains from mice with either a *Tgfbr2^fl/wt^* or *Tgfb1^fl/wt(ex1)^* allele already in our colonies. In neonatally-induced *Cx3cr1^CreER(Litt)^* mice, we observed wide-spread loss of the homeostatic marker P2ry12 in IBA1^+^ microglia, accompanied by their adoption of an activated morphology (Fig 6a). In contrast, and similar to the published report^27^, tamoxifen-administration to *Cx3cr1^CreER(Litt)^* mice after P21 had no apparent effect on microglia homeostasis or activation (Fig 6b). Likewise, we observed no apparent phenotypic changes in neonatally-induced *Cx3cr1^CreER(Jung)^* or *P2ry12^CreER^* mice. These experiments were repeated and confirmed in two different research laboratories (Luo Lab, University of Cinncinnati; Arnold Lab, University of California, San Francisco) to control for the possibility that differences in vivarium microbiomes or research technique (e.g. IP injection-induced peritoneal infection/inflammation), as was recently described in cerebral cavernous malformation (CCM) mouse models^28^, might contribute to the observed loss of microglia homeostasis in *Cx3cr1^CreER(Litt)^* mice.

**Figure 6.**
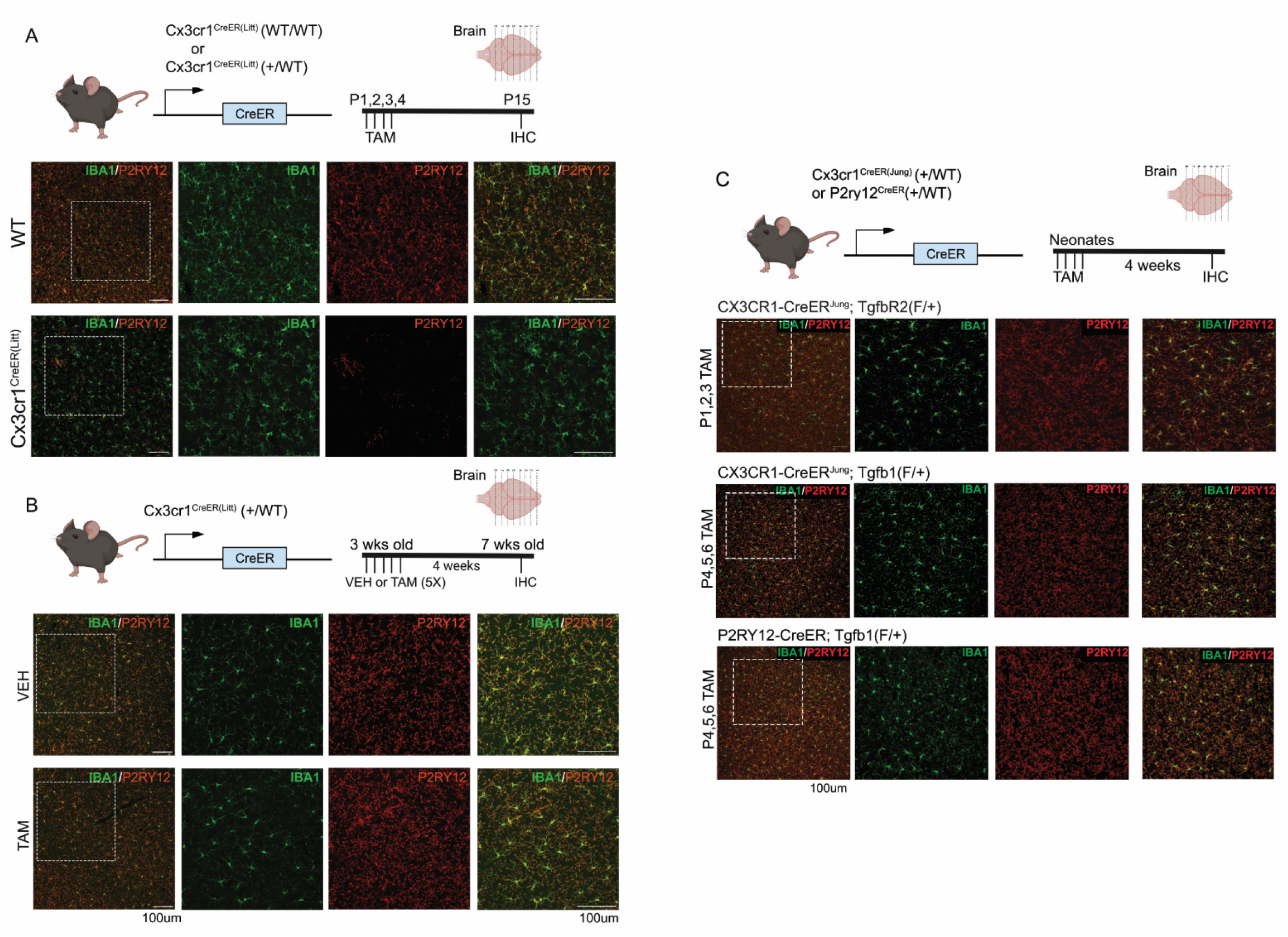
Evaluation of the dyshomeostatic microglia in different icroglia-specific CreER drivers after TAM treatment at different ages. P2Ry12 expression is used as a measure of dyshomeostasis in microglia. (A), consistent with previous studies, we observe dyshomeostasis of microglia (indicated by loss of P2RY12 expression) across many region in the neonatal Cx3cr1^CreER(Litt)^ (+/WT) mice treated with TAM. This phenotype is not observed in (B) the adolescent (3wk old) Cx3cr1^CreER(Litt)^ (+/WT) mice that received TAM treatment or (C) neonatal Cx3cr1^CreER(Jung)^ (+/WT) and P2RY12CreER (+/WT) mice that received TAM at the similar time frame. Scale bar= 100µm.

In investigating the microglia phenotype observed in *Cx3cr1^CreER(Litt)^* mice, we were struck by its similarity to microglia deficient in TGFβ-signaling (*Itgb8;nestinCre and Tgfbr2^fl/fl^;Cx3cr1^CreER^ ^29^, Tgfb1^-/-30^; Tgfbr2^fl/fl^Sall1^CreER^ ^31^; Lrrc33/Nrros^-/-15^*), including loss of homeostatic signature genes such as *P2ry12*, *HexB*, *SiglecH* and *Tmem119* with reciprocal upregulation of MgND/DAM (neurodegeneration-associated phenotype by microglia/disease-associated microglia) signature genes including ApoE, Axl, and Lglas3; and genes directly or indirectly related to interferon signaling including Irf7, Siglec1, and Mx1 (Fig 6a). Indeed, the transcriptional phenotype of microglia from neonatal (P15) *Tgfbr2^fl/fl^;Cx3cr1^CreER^*mice are moderately correlated to those from neonatal tamoxifen-treated Cx3cr1^CreER(Litt)^ mice (R=0.56; P>2e-16). Importantly, these various TGF-β-signaling deficient mouse models were not generated using the *Cx3cr1^CreER(Litt)^* line, raising the hypothesis that neonatal tamoxifen administration to *Cx3cr1^CreER(Litt)^* mice might somehow dysregulate TGF-β signaling. We directly tested this by immunostaining mouse brains for phosphorylated (p)Smad3, an indicator of canonical (SMAD-mediated) TGF-β signaling. While there was a trend for reduced pSmad3 immuno-fluorescent staining in microglia from TAM-treated compared to non-TAM treated controls, the overall change on the population level was not significantly different. These results suggest that reduced TGF-β1-signaling is not likely a primary driver of the microglial phenotype in *Cx3cr1^CreER(Litt)^*mice, but could be a downstream consequence of increased INF-signaling as proposed by Suhasrabuddhe et al. Consistent with this, INF and TGF-β1/Smad signaling are co-regulated in activated microglia^32^, and Smad2/3 is known to directly regulate the expression of several homeostatic markers including P2ry12^33^.

## Discussion

The use of the Cre-LoxP system has revolutionized biological research by enabling cell-type-specific gene manipulation. The addition of temporal control to Cre activity was introduced by the addition of a modified estrogen receptor (ER) ligand binding domain^34^ providing exquisite precision in targeting and tracking ontogenically distinct cell subtypes and their associated lineages. The application of Cre-based genetic targeting to microglia biology has had particularly important impacts on our understanding of brain immunity. Over the past decade, research utilizing two *Cx3Cr1^CreER^* lines ^16,35^ helped to elucidate the ontogeny of microglia and non-microglia CNS-associated macrophages, and the roles of specific microglia genes and signaling pathways in brain development and function^35,36^. Recently, in an effort to improve the specificity of brain microglia gene manipulation, several new brain microglia-specific inducible CreER mouse lines were generated, including *Tmem119^CreER^, P2ry12^CreER^ Sall1^CreER^, HexB^CreER^* ^8,10,31,37^. Since the initial reports characterizing these new lines, we and others noticed different degrees of leakiness and recombination efficiency. Furthermore, recent reports document important off-target effects in *Cx3cr1^CreER(Litt)^* mice^12,27^, prompting us to perform a more detailed characterization of the publicly-available microglia-targeting CreER lines. Our studies focus on five key characteristics of CreER gene targeting: specificity, leakiness, efficiency, extra-neural recombination, and off target effects. Because *Sall1^CreER^* and *Hexb^CreER^* were not readily available to us at the time of this report, we compare our results to published studies using these lines^31,37^. We believe this evaluation will provide valuable information to the field, particularly for researchers aiming to identify appropriate mouse models for conditional gene deletion in microglia.

**Table 1:** Summary of key features across different microglia CreER driver lines.

|  |  |  |  |  |  |  |  |  |
| --- | --- | --- | --- | --- | --- | --- | --- | --- |
| Mouse Line | Specificity | Leakiness | Efficiency | Extraneural Recombination |  | Off-Target Effects | Use Information |  |
|  |  |  |  | Blood | Spleen |  | Availability | Reference |
| CX3CR1<sup>CreER</sup>Litt | + | +++ | +++ | ++ | +++ | +++ | JAX: 021160 | Parkhurst et al., 2013 |
| Cx3cr1<sup>CreER</sup>Jung | + | ++ | ++ | ++ | ++ | + | JAX: 020940 | Yona et al., 2013 |
| P2ry12<sup>CreER</sup> | ++ | + | + | - | + | - | JAX: 034727 | McKinsey et al. 2020 |
| Tmem119<sup>CreER</sup> | ++ | + | + | - | - | - | JAX: 031820 | Kaiser et al., 2019 |
| HexB<sup>CreER</sup> | +++ | + | + | +++ | ++ | + | Not Publicly available | Masuda et al., 2020 |
| Sall1<sup>CreER</sup> | + | ++ | + | - | ? | + | Not Publicly available | Buttgereit et al., 2016 |

### Specificity

*Cx3cr1^CreER^* (Littman and Jung) recombine microglia and other CNS (brain border) macrophages, myeloid precursors, and dendritic cell subsets in the meninges. *P2ry12^CreER^* specifically recombines microglia, in addition to a small subset of dural and choroid plexus macrophages (likely epiplexus cells)*. Tmem119^CreER^* recombines microglia and non-macrophage meningeal/perivascular cells (likely fibroblasts). *HexB^CreER^* was shown to provide specific CNS-microglia recombination without labeling neurons, astrocytes, oligodendrocytes or vascular cells in the brain, and can distinguish CNS associated macrophages (CAMs) from microglia. *Sall1^CreER^* recombines microglia specifically, in addition to neurons, astrocytes, and oligodendrocytes.

### Leakiness

We found that *Cx3cr1^CreER^* lines recombined the more sensitive Ai9 tdTomato reporter a relatively large number of microglia (and splenic macrophages) in the absence of tamoxifen, whereas *Tmem119^CreER^*and *P2ry12^CreER^* lines were comparatively less leaky. *Sall1^CreER^*is relatively more leaky than *Hexb^CreER^*, and similar to *Cx3cr1^CreER(Jung)^*. These three lines were directly compared using both *RosaYFP* and *Ai9* reporter mice to assess leak^37^. Like us, this group observed that leakiness was negatively correlated with recombination efficiency or the length of the loxP flanked region (higher leakiness in *Ai9* reporter and lower leakiness in *R26-YFP* reporter).

### Efficiency

Efficiency was directly correlated with inter-loxP distance and intrinsic CreER activity/expression. *Tmem119^CreER^* and *P2ry12^CreER^*mice with the *Ai14* reporter (0.9kb) maintained high recombination efficiency but only achieved partial recombination in the *R26-YFP* reporter (2.7kb) allele (about 30% of all microglia population), reiterating the importance of loxP distance in recombination efficiency. In line with this conclusion, we found that the *Cx3Cr1^CreER(Jung)^* line effectively deletes both copies of the shorter Tgfb1^fl/fl^ target gene and the longer Alk5^fl/fl^ gene while the *P2ry12^CreER^*had a 50% decrease in mRNA levels of these two genes. It is worth noting that this 50% gene deletion efficiency in the *P2ry12^CreER^* line was evaluated in sorted YFP+ microglia populations. As *P2ry12^CreER^* only achieves *R26-YFP* recombination in about 30% of total microglia, the gene deletion efficiency in the full population of microglia sorted using the *Ai9* tdTomato reporter was lower (20% decrease in Alk5 mRNA levels in total tdT^+^ cells). Masuda et. al^37^directly compared *HexB^CreER^*, *Sall1^CreER^* and *Cx3cr1^CreER(Jung)^*, and found *HexB^CreER^* and *Sall1^CreER^* to be similarly less efficient than *Cx3cr1^CreER^*. Note that most of their analysis was done using homozygous *Hexb^CreER/CreER^* mice which likely increased recombination efficiency, as we observed with P2ry12^CreER/CreER^ mice.

### Extraneural recombination

Outside of the CNS, *Cx3cr1^CreER^* (Littman and Jung) recombine peripheral blood and peripheral immune cells including dendritic cell subsets, T cells, NK cells, and tissue-macrophages in most organs including the spleen. *P2ry12^CreER^* recombines a small subset of tissue-resident macrophages including a population of splenic macrophages. *P2ry12^CreER^* does not recombine blood cells or platelets. *Tmem119^CreER^* recombines a small percentage of IBA1^+^ macrophages and a substantial number of IBA1-negative cells in the spleen (the exact cell type of these tdTomato+ cells are not clear). In comparison, *Sall1^CreER^* does not recombine blood cells, while *HexB^CreER^* mice have significant acute recombination in Ly6c^hi^ and Ly6c^lo^ cells (similar to *Cx3cr1^CreER(Jung)^*line), and additionally recombine nearly all circulating Ly6G^+^ granulocytes^37^ (Supplementary Figure 11). Recombination of blood cells by *Hexb^CreER^*is paralleled by tdT expression in most F4/80+/Iba1+ splenic myeloid cells^37^ (Supplementary Figure 6) from HexB^tdt/tdt^ reporter mice.

### Off target effects

Our study explored a phenomenon described by Sahasrabuddhe and Ghosh 2022. This group found that neonatal TAM administration to the *CX3CR1^CreER(Litt)^* line resulted in DNA damage, induction of microglial interferon-1 signaling and an altered microglia transcriptional profile. (This effect was not observed in the adult tamoxifen administered *CX3CR1^CreER(Litt)^* mice, or in *Tmem119^CreER^*mice with neonatal TAM administration.) Interestingly, the transcriptional phenotype of microglia from neonatally-induced *Cx3cr1^CreER^*^(Litt)^ is similar to *Sall1flox/Sall1CreER, Sall1flox/flox;Cx3cr1^CreER(Jung^*^)^ and Tgfbr2^fl/fl^(*Cx3cr1^CreER^*) with increased expression of INF-response genes and other DAM/MgND genes, and downregulation major homeostatic microglia genes including *P2ry12, Tmem119, HexB* and *Sall1* itself. This similarity prompted us to explore whether TGF-β/Smad signaling might directly be affected by neonatal tamoxifen induction in *Cx3cr1^creER(Litt),^* or other microglia-targeting Cre lines. We found that despite robust activation and loss of P2RY12 expression in microglia in *Cx3cr1^CreER(Litt)^* mice, pSMAD3 immunofluorescence intensity (a measure of TGF-β signaling) was not appreciably changed. Furthermore, we observed no evidence of alterations in microglia homeostatic gene expression in *Cx3cr1^CreER(Jung)^* and *P2ry12^CreER^* lines.

We and others previously investigated whether the gene knock-in strategies used to generate these various CreER lines might affect gene expression resulting in unintended effects. *Cx3cr1^CreER^* (Littman and Jung) and Sall1^CreER^ lines were generated using a knock-in/knock-out strategy and so by definition have reduced expression of Cx3cr1 and Sall1, respectively. Both of these genes are known to have important functions in brain development. Tmem119-, P2ry12- and Hexb-CreER lines were generated CRISPR-facilitated homologous recombination in which the target gene stop codon was replaced by a ribosome skipping fusion peptide coding sequence, followed by the coding sequence for CreER. All lines show some diminution in mRNA expression of target gene expression. No brain-specific functions have been found for Tmem119. P2RY12 signaling is responsible for inflammation-induced microglia process exptension^38^ and microglia containment of leakage following cerebrovascular injury^39^. Homozygous loss of *HexB* in mice causes a fatal demyelinating microgliopathy similar to *HexB* inactivating mutations in patients with Sandhoff’s Lysosomal Storage Disease. In light of the known roles for Cx3cr1, Sall1, P2ry12, and Hexb in development and disease, it is at least theoretically possible that reduction in their gene expression could have unintended experimental effects.

### Additional considerations and recommendations

While the *Cx3Cr1^CreER(Litt)^* and *Cx3Cr1^CreER(Jung)^*lines have been used widely in the field, alternative microglia *CreER* lines are still new to the field. This research highlights the value of a systematic comparison and evaluation of genetic tools to better design experimental strategies for microglia conditional gene manipulation. While we tried to be comprehensive in our assessment of the currently available tools, there are still several microglia-specific lines that we did not investigate, including *Sal1CreER* and *Hexb-CreER,* and a new Fcrls-2A-Cre available in JAX (036591). Based on our results, we are able to make the following recommendations regarding choice of CreER line and specific experimental goals.

If the key question being investigated requires deletion in all microglia, the two *Cx3cr1^CreER^* lines are advantageous due to their high recombination efficiency. Comparing the two *Cx3Cr1^CreER^* lines (Littman and Jung), our data suggest that the *Cx3^CreER(Jung)^* line has less Cre activity in the absence of TAM (both with the *Ai9* and *R26-YFP* reporter) while having almost equally high recombination efficiency after TAM treatment. The fused IRES-eYFP reporter in the *Cx3Cr1^CreER(Littman)^* is convenient for sorting microglia without the need for surface antibody staining such as CD11b and CD45. However, crossing the *Cx3Cr1^CreER(Jung)^* line with the *AitdTomato* or *R26-YFP* reporter lines can be easily accomplished to facilitate genetic reporter-based FACS sorting. As peripheral macrophages also express *Cx3Cr1* and are therefore recombined by *Cx3cr1^CreER^*, peripheral immune effects are a potential confound that needs to be considered when using these two lines.

To investigate gene function in “true” microglia, *Tmem119^CreER^*, *P2ry12^CreER^* are preferable. Additional consideration should be taken with *Hexb^CreER^* which was found to strongly recombine blood cells. Utilization of the *Tmem119^CreER^* or *P2ry12^CreER^* lines therefore enables enriched labeling of CNS parenchyma microglia and allows researchers to track endogenous microglia and study their specific properties after CNS injury or in neurodegenerative conditions. This was successfully achieved in an EAE model using the *P2ry12^CreER^* line^19^. Future use of these tools could facilitate the pre-labeling of brain microglia cells and allow for the separation of microglia from infiltrating peripheral monocytes/macrophage by FACS sorting. This will help resolve a long-standing question in the field: whether the infiltrating monocytes/macrophage have a distinct neuroinflammatory program compared to the resident brain microglia in CNS disease or injury. The lower recombination efficiency observed in *P2ry12^CreER^* and *Tmem119^CreER^* also lends itself well to performing mosaic genetic experiments which allow for *in vivo* single-cell phenotypic analyses, where differences among single cells can be attributed to induced gene mutation or expression in an otherwise identical organism and genetic background.

We attempted to enhance recombination efficiency of the *P2ry12^CreER^* line by crossing to the iSuReCre reporter mouse. Compared to *DCX^CreER^*;iSuReCre mice, with robust recombination of immature DCX+ neuroblasts, we observed no microglial recombination in *P2ry12CreER*;iSuReCre mice based on immunostaining brain sections for the iSuReCre membranous tomato report. Interestingly, iSuReCre shows reliable recombination and tdTomato expression in peripheral macrophages by LysMCre, and Tie2Cre;iSuReCre mice have apparently strong recombination of retinal microglia. The underlying cause for lack of iSuReCre recombination by *P2ry12^CreER^* mice is unknown but may be due to the genomic integration site of the iSuReCre-tdT construct which has not been as well-characterized as the ROSA26 safe-harbor locus. Our data in the *Cx3cr1^CreER(Jung)^*-iSureCre mice show that even with the strong *Cx3cr^CreER^*driver, MbTomato expression is absent in microglia (Data not shown), therefore supporting the hypothesis that the specific loci where iSuRe-Cre cassette is inserted might be silenced in microglia. An alternative method to boost CreER activity is to breed the line to homozygosity. *P2ry12^CreER/CreER^* (unpublished observations) and *Hexb^CreER/CreER^* mice have more efficient gene deletion. However, this may come at the cost of increased leakiness and potential off-target effects related to gene dosage. Additionally, an investigator can choose to breed in one knock-out allele to reduce the number of recombination events required for complete gene deletion.

When using the *Tmem119^CreER^* or the *P2ry12^CreER^* lines, we recommend using a reporter with inter-loxP distance as close to that of the target gene as possible, to include one knockout allele so that only one recombination event is required for gene deletion (if heterozygous knockout mice do not cause phenotype), or ideally to utilize one target allele that is floxed with an intrinsic reporter (knock-in/knock-out construct) and one knockout allele to simultaneously facilitate gene deletion and precisely identify knockout cells. Understanding that null and conditional knockout-knockout reporters, and other Cre “boosting” constructs, are not readily available, additional analysis such as qRT-PCR on sorted microglia, RNAscope or immunohistochemical staining is recommended to validate successful knockout or knockdown of the target genes in brain microglia. We recommend performing this validation in a small pilot study before investing time and resources in larger scale experiments.

With our rigorous and comprehensive analysis of the four commercially available microglia-CreER mouse lines on the five key features we chose to focus on, we identify important caveats and strengths for these lines which will be of broad significance for researchers interested in performing conditional gene deletion in microglia. While we are preparing for this manuscript, a preprint of an independent study evaluating microglial CreER lines was reported^40^, including evaluation of *HexB^CreER^* mice. Our two studies provide both overlapping and distinct recommendations and guidelines for the use of these powerful tools. Together with this study, we hope our comprehensive comparison and analysis provides important data and guidance for researchers interested in performing conditional gene manipulation in microglia.

**Figure 7.**
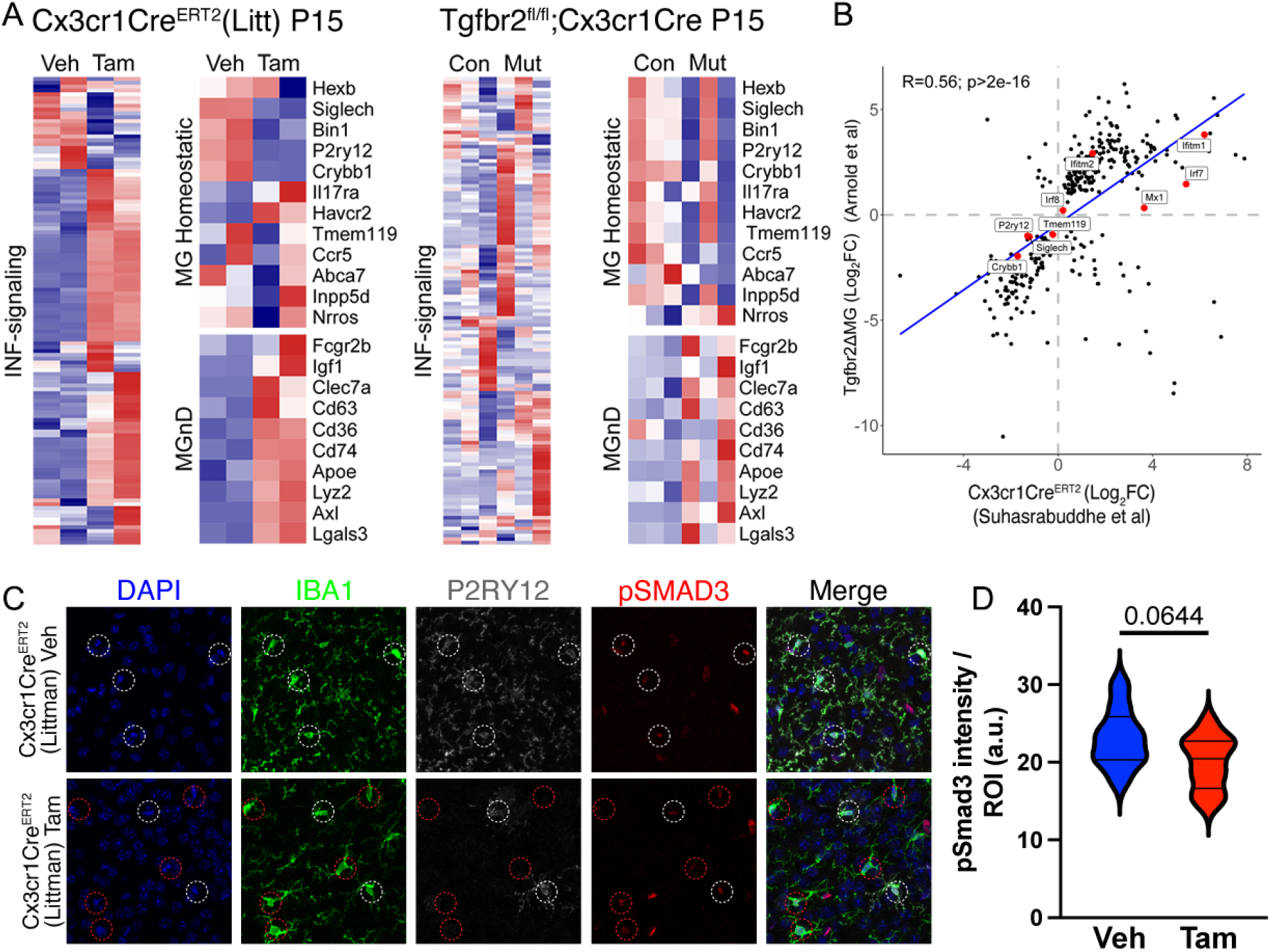
Similarity in microglia phenotypes from neonatal tamoxifen-induced Cx3cr1CreER(Litt) and Tgfbr2fl/fl;Cx3cr1Cre mice. (A) Transcriptomic phenotypes from P15 Cx3cr1CreER(Litt) (vehicle versus tamoxifen) and Tgfbr2;Cx3cr1Cre (Tgfbr2fl/+;Cre vs Tgfbr2fl/fl;Cre) mice showing changes in INF-signaling related, MgND/DAM, and hemeostatic gene expression. (B) Correlation between two data sets. (C) Brain sections from P15 Cx3cr1CreER(Litt) (vehicle versus tamoxifen) mice stained for DAPI (blue), IBA1 (green), P2ry12 (grey), pSMAD3 (red). Quantification of pSMAD3 mean fluorescence intensity in individual microglia from each group (D) shows no significant difference in pSMAD3 (P=0.0644; Students T-Test). Data were combined from 3 vehicle-treated and 4 tamoxifen-treated mice. Scale bar: 100 μm.

## Acknowledgement

Y.L is supported by NIH grants (R01NS127074, R01NS107365, R21NS127177). A.B is supported by NIH 1F31NS125930-01. K.W is supported by NIH 1F31NS129204-01A1. T.A is supported by NIH grants (1R01NS119615-01;R01NS123168). N.S is supported by institutional funds. We thank Chet Closson and the University of Cincinnati live imaging core (supported by NIH S10OD030402) for technical support. We would like to thank Dr. Rui Benedito for sharing the iSuReCre mouse line and Dr. Dr. Zhi-qi Xiong for sharing the *DCX^CreER^* mouse line.

## Author contributions

YL and AB conceptualized the study. AB, GM, TA and YL designed the experiments. AB performed most of the in vitro and in vivo experiments, recorded and analyzed data with help from KW. GM carried out part of the neonatal TAM experiment and NS performed the bioinformatic data analysis. AB, GM, TA and YL drafted and revised the paper. All authors read, edited, and approved the final version of the manuscript.

## Supplementary material is available online

**Table S1.** Quantification of the FACS analysis of the reporter+ microglia in whole brain single cell suspension prepared from the P2RY12CreER-Ai9-R26-YFP mice after TAM treatment. Related to Fig 2

## STAR Methods

### RESOURCE AVAILABILITY

#### Lead contact

Further information and requests for resources and reagents should be directed to and will be fulfilled by the Lead Contact, Yu Luo.

#### Materials availability

This study did not generate new unique reagents.

#### Data and code availability

- We analyzed recently published publicly available RNA-seq data sets: GEO: GSE190207 and GEO: GSE124868
- Microscopy data and behavioral test data reported in this paper will be shared by the lead contact upon request.
- No original code was generated in this study.
- Any additional information required to reanalyze the data reported in this paper is available from the lead contact upon request.

### EXPERIMENTAL MODEL AND SUBJECT DETAILS

#### Animals

All animal protocols were approved by the IACUC of University of Cincinnati or the IACUC of University of California, San Francisco. All transgenic lines (see details below) and C57BL/6J WT mice were purchased from Jackson Laboratory and housed in the animal facility of the University of Cincinnati or University of California, San Francisco. Mice were maintained with a 12-hour light/dark cycle and fed ad libitum. To evaluate cre recombinase specificity and efficiency, we utilized four different microglia CreER lines: *Cx3cr1^GFPCreER(Litt)^*Line (JAX stock number 021160)^41^; *Cx3cr1^CreER(Jung^*^)^ Line (JAX stock number 020940)^35^; *Tmem119^CreER^* line (JAX stock number 031820)^42^ and the *P2ry12^CreER^*line (JAX stock number 034727)^19^. For fluorescent reporter lines, we crossed the above microglia CreER lines with either the *Ai9 R26-CAG-tdTomato* line (JAX stock number 007909)^43^ or the *ROSA26-YFP* line (JAX stock number 006148)^44^ or both of the reporter lines in some of the experiments. To analyze target gene deletion efficiency with the different floxed l between the two loxP sites, we generated *Cx3cr1^CreER(Jung)^*or *P2ry12^CreER^*-*Tgfb1^fl/fl^* mice (floxed exon 3, ∼500 bp, JAX stock number 65809) or *Cx3cr1^CreER(Jung)^* or *P2ry12^CreER^-Alk5^fl/f^*^l^ mice (Floxed region is between 1.6-1.7 kb, JAX stock number 028701)^45^. To investigate whether the iSuRe-Cre transgene is inducible in microglia in the *P2ry12^CreER^* line and the *DCX^CreER^* line^46^, iSuRe-Cre^20^ mouse line is obtained from Dr. Rui Benetido (Centro Nacional de Investigaciones Cardiovasculares -CNIC, Spain) and crossed with the *P2ry12^CreER^* or *DCX^CreER^*mouse line to generate *P2ry12^CreER(+/WT)^*iSuRe-Cre^(+/WT)^ or *DCX^CreER(+/WT)^*iSuRe-Cre ^(+/WT)^ mice. No animals were excluded from data analysis except where tissue preservation quality was not ideal due to unsuccessful perfusion (indicated by shredded tissue) or poor cryoprotection in tissues.

#### Tamoxifen treatment in vivo

The variety of MG^CreER^-Ai9 or MG^CreER^-*R26YFP* or *P2ry12^CreER^-Ai9/R26YFP* double reporter mice and the TGF-β1 and ALK5 floxed mice (8-12 weeks old, both male and females) were given tamoxifen (TAM) dissolved in 10% EtOH/90% sunflower oil or vehicle without TAM by gavage feeding at a dose of 180 mg/kg daily for 5 consecutive days. This dosing regimen was previously demonstrated to provide maximal recombination with minimal mortality and successfully monitored the adult NSCs in previous studies by our group and the others^47,48^. To evaluate recombination efficiency in the brain and the clearance of splenic monocytes/macrophages, we collected the brain and the spleen of VEH or TAM treated mice at 4 weeks after the treatment for immunohistochemical (IHC) or flow cytometry (FACS) analysis. For studying the phenotype of non-homeostatic microglia in CreER(+/WT) mice with TAM treatment, *Cx3cr1-^CreER(Litt)(+/WT)^*neonatal mice were subjected to either VEH or TAM treatment at neonatal days of P1-P4 (50µg via intragastric injection) and brain harvested for analysis at p15. To test effect of TAM treatment in the *Cx3cr1-^CreER(Litt)(+/WT)^* adolescent mice, mice at 3 weeks of age were treated with 5 day gavage (180mg/kg daily) and harvested at 4 weeks after TAM treatment. Neonatal *Cx3cr1^CreER(Jung)^* allele carrying mice (*Cx3cr1^CreER(Litt)(+/WT)^tgfbr2^wt/fl^ or tgfb1^WT/fl^*) or *P2ry12^CreER^* allele carrying mice (P2ry12^CreER+/WT^tgfb1^WT/fl^) were subjected to TAM treatment at neonatal days of P1-P4 (50µg via intragastric injection) and brain harvested for analysis at P15 or at neonatal days P4-P6 (500ug via intraperitoneal injection) and harvested at P30.

### METHOD DETAILS

#### Immunohistochemistry

Mice were anesthetized and perfused with PBS or PBS followed by 4% paraformaldehyde (PFA). For brains that were subjected to both IHC and FACS analysis, mice were only perfused with PBS and part of the brain block was drop fixed in 4% PFA overnight before being transferred to 20% then 30% sucrose. For mice that were perfused by 4% PFA, the brain and spleen was dissected and post-fixed in 4% PFA overnight at 4 °C and equilibrated in 20% then 30% sucrose. 30 μm-thick sections were cut in a Leica Cryostat and blocked in 4% BSA/0.3% Triton-x100 for 1 hour. After blocking, sections were incubated with primary antibodies for 18h-42h at 4 °C and followed by appropriate secondary antibodies conjugated with Alexa fluorescence 488, 555, 647 or 790. The following primary antibodies were used in this study: IBA1 (Rabbit 1:1000 Wako), P2RY12 (Rabbit 1:200 Anaspec), P2RY12 (Rat 1:500 BioLegend), GFP (Rabbit 1:1000 Invitrogen), PSMAD3 (Rabbit 1:100 Abcam), NEUN (Mouse 1:1000 BioLegend), DCX (Rabbit 1:1000 Cell Signaling). Omission of primary or secondary antibodies resulted in no staining and served as negative controls. Images were acquired by a motorized stage-equipped Leica DM5000B microscope (Leica Microsystems, Bannockburn, IL) equipped with Stereo Investigator image software (MBF Bioscience, Williston, VT) for unbiased sampling of the tissue field. Confocal imaging was carried out on a Leica Stellaris 8 confocal microscope (Leica Microsystems, Bannockburn, IL). Random imaging field was selected by SI with outlined ROI and grid and images were taken at 20X or 40X objectives. Quantification of the percentage of reporter positive cells (tdTomato+ or YFP+) among IBA1+ microglia (in the brain) or macrophage (in the spleen) was carried out by as described in our previous publication^49^ using Nikon Element or ImageJ software. At least 3 sections containing the region of interest at similar coronal locations were quantified for each mouse and averaged values (total or average) for each animal were considered as one data point for statistical analysis. Group and treatment information were all blinded to the image analyzer.

#### FACS sorting

Fluorescent activated cell sort (FACS) was accomplished using a papain dissociation system (9001-73-4, Worthington Biochemical Corporation). To collect tissue, animals were perfused with cold 1x HBSS for 2-3 minutes. The brain was then extracted and mechanically dissociated with a scalpel before using the papain dissociation kit. Once dissociated, to remove excess myelin and debris cells were suspended in a 37% percoll solution and spun at 800g for 20 minutes. The cells were then resuspended in FACS buffer (0.5% BSA in PBS) for sorting. Gating was determined using the tdTomato or yellow fluorescent protein reporter (YFP) expressed by microglia cre line compared to cells isolated from WT control mice for microglia collection on a BD/FACSAria II (BD Biosciences, Franklin Lakes, NJ). For analysis of targeted floxed allele expression (*Tgfb1^fl/fl^*or *Alk5^fl/fl^*) in a variety of microglia-specific CreER lines, we sorted YFP+ or tdTomato cells at 3-4 weeks after TAM administration and processed the sorted microglia cells for qRT-PCR for gene expression analysis as described in details below.

#### qRT-PCR analysis to evaluate reduction of mRNA levels of the floxed gene alleles (Tgfb1 and Alk5)

RNA expression levels for housekeeping genes (Hmbs, PGK1), Iba1 and targeted floxed genes (*Tgfb1^fl/fl^* or *Alk5^fl/fl^*) were determined by Reverse-Transcribed quantitative real-time PCR. RNA was extracted from FACS sorted YFP+ or tdTomato+ cells using a RNAqueous-Micro Total RNA isolation kit (AM1931, ThermoFisher Scientific). Total RNA was treated with RNase free DNase and cDNA was then generated using iScript cDNA synthesis kit (1708890, BioRad). cDNA levels for Hmbs (hydroxymethylbilane synthase), pGK1 (phosphoglycerate kinase 1) and various target genes were determined, using specific primer/probe sets by quantitative RT-PCR using a Roche Light Cycler II 480 (Roche, Basel, Switzerland). Relative expression level was calculated using the delta Ct method compared to Hmbs as a reference gene and expressed as fold change compared to the average of WT cells for each individual gene. Primers and carboxyfluorescein (FAM) labeled probes used in the quantitative RT-PCR for each gene are listed in the oligonucleotide section. To selectively detect the presence of the floxed region in the TGFb1^fl/fl^ or the ALK5^fl/fl^ microglia, the probe based qRT-PCR reactions were selected so that the forward and reverse primers span the exon junctions of the floxed exon and a neighboring exon therefore ensuring the loss of amplification in the case of a successful cre-lox mediated recombination.

#### Bioinformatics analysis and quantification of pSMAD3 immunoreactivity in microglia

We reanalyzed datasets from GSE190207 (*Cx3cr1^CreER^*Littman) and GSE124868 (*Tgbfr2 fl/fl;Cx3cr1^Cre^* at P15). Fastq files were aligned to the mouse genome (mm10) with Rsubread 2.10 and quantified using FeatureCounts. Differential expression analysis was performed with DESeq2 1.36. Heatmaps were made with the pheatmap 1.0 package, while scatter plots were made with ggplot2 and ggrepel.

Quantification of pSMAD3 immunofluorescent staining was determined as previously published^29^ using stained cryosections from four neonatal TAM-treated *Cx3cr1^CreER(Litt)^* and three vehicle controls at P15. Three randomly chosen confocal images from each sample were taken using the same confocal settings. ImageJ software was used to quantify the number of microglia cell nuclei (DAPI-positive, Iba1-positive), and the intensity of pSMAD3 staining in each microglia cell.

### QUANTIFICATION AND STATISTICAL ANALYSIS

All studies were analyzed using SigmaPlot. Results are expressed by mean±SEM of the indicated number of experiments. Statistical analysis was performed using the Student’s t-test, and one- or two-way analysis of variance (ANOVA), as appropriate, with Tukey post hoc tests. A p-value equal to or less than 0.05 was considered significant.

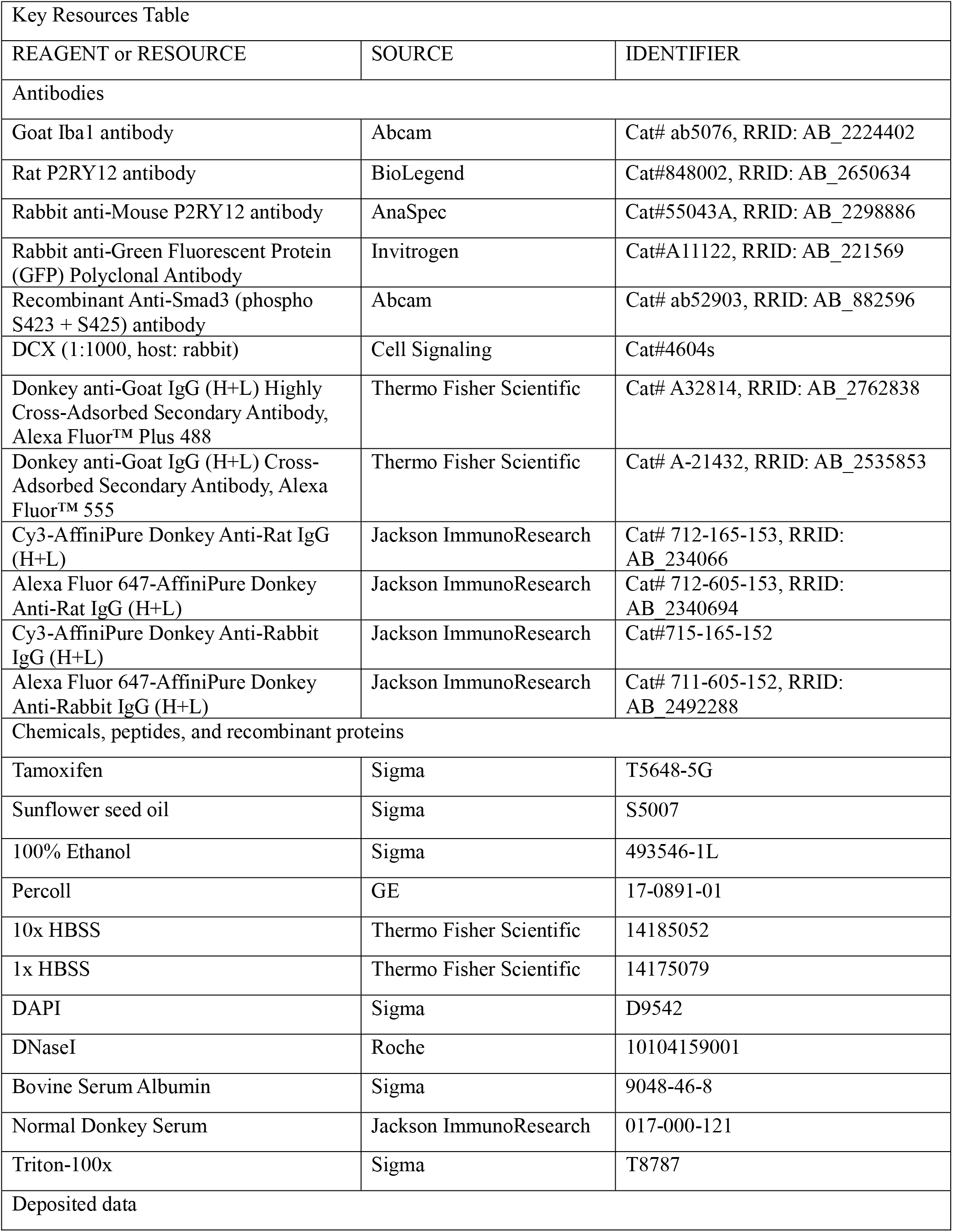
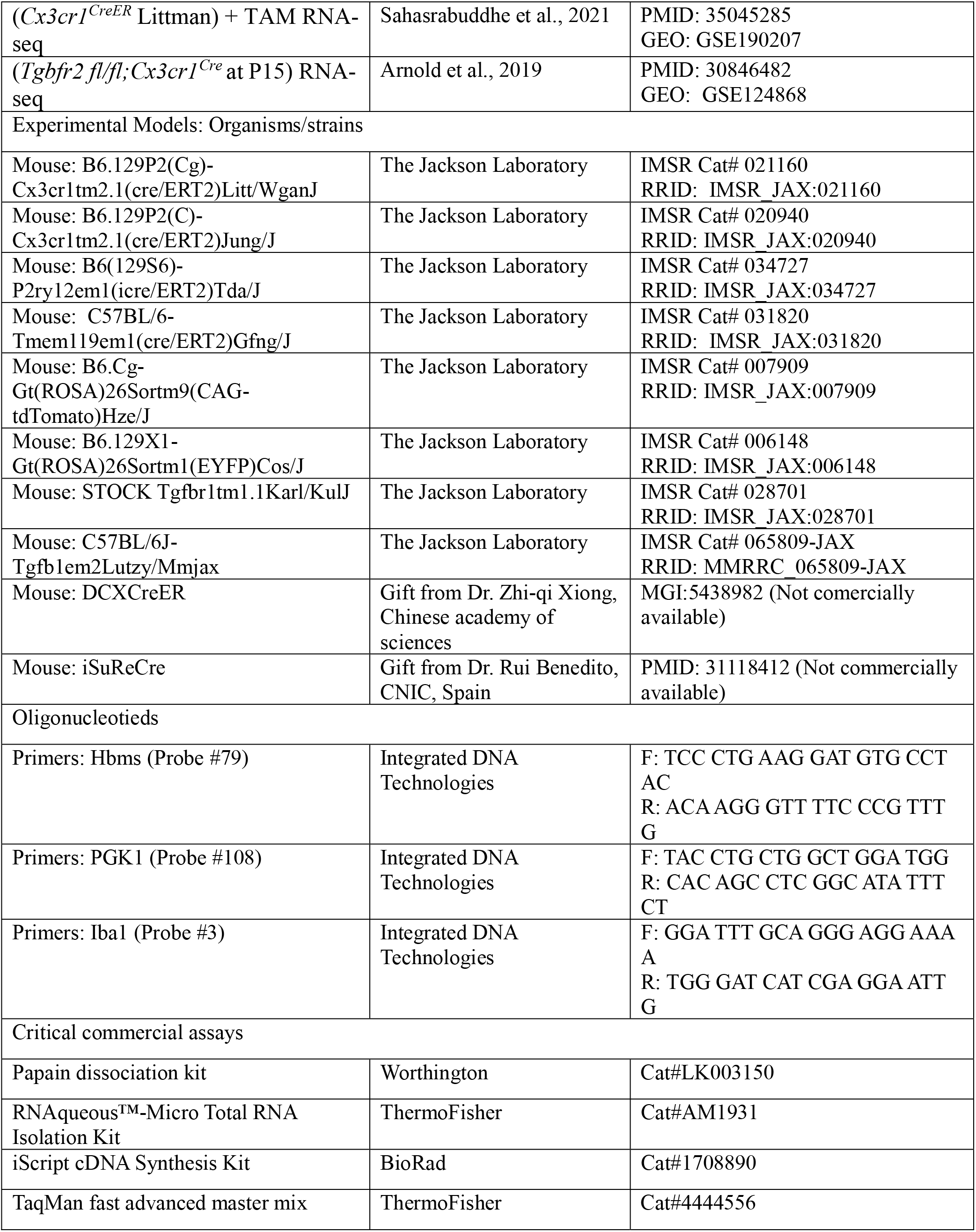
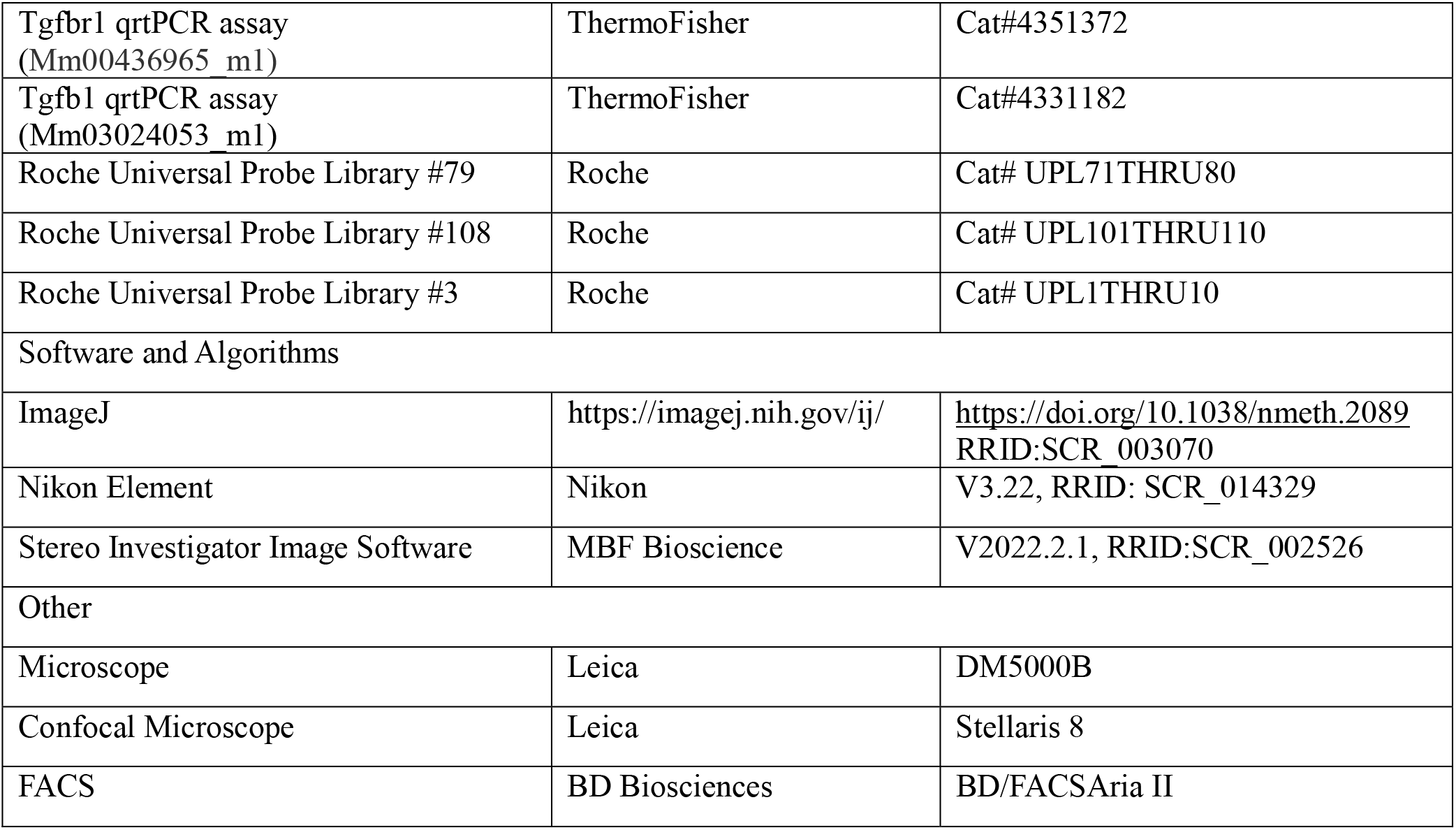

---

**Supplementary Figure 1.**
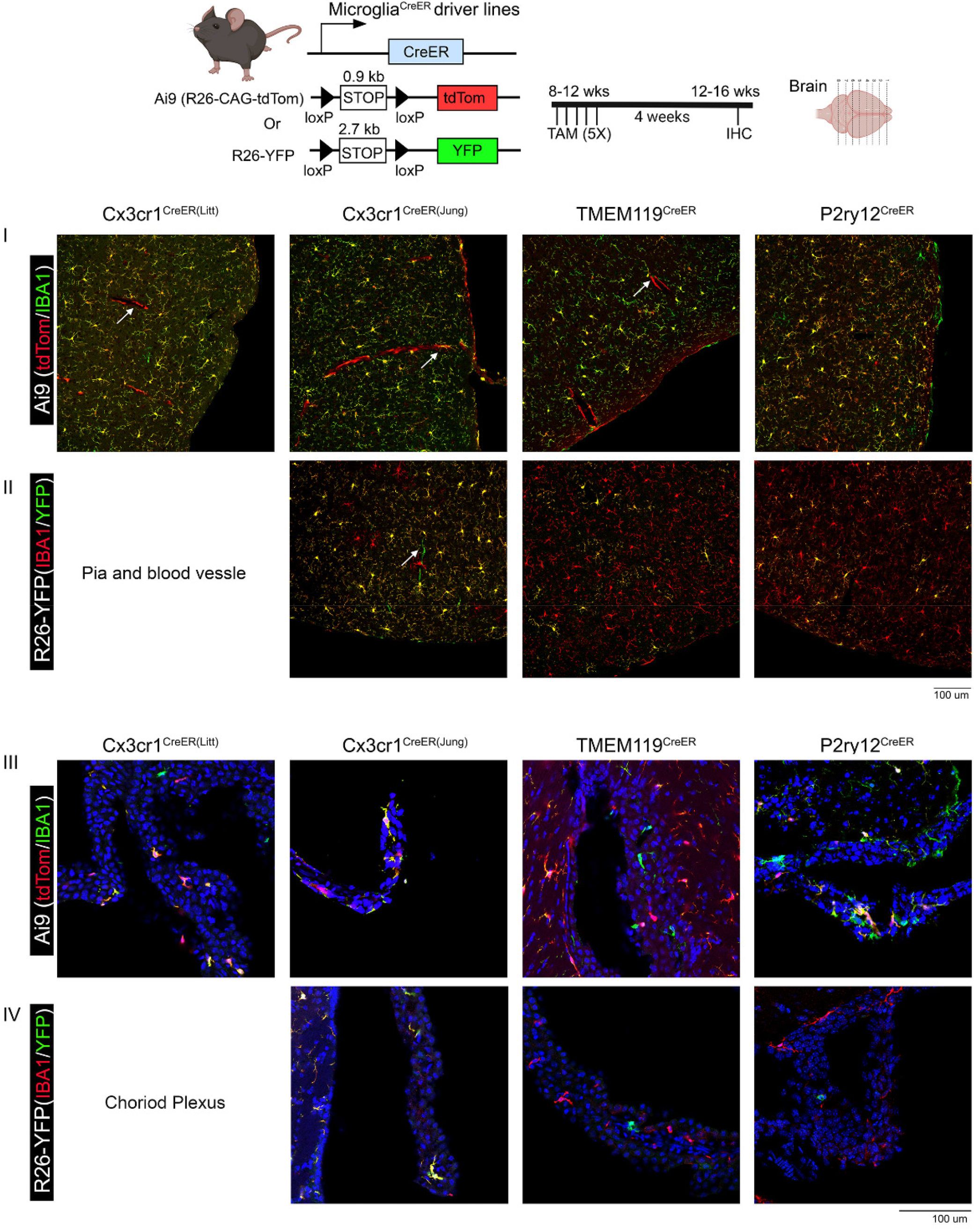
Border Associated macrophage (BAMs) labeling in the four different creER driver lines using either the Ai9 (tdTomato) or R26-YFP reporter mouse lines. The experimental timeline is shown on top of the panel. Representative images from each cre driver and Ai 9 (I) or R26-YFP (II) reporter line in pia and blood vessel associated macrophages and choroid plexus associated macrophages (III Ai tdTomato) or (IV R26-YFP) reporter. Scale bar = 100μm.

**Supplementary Figure 2.**
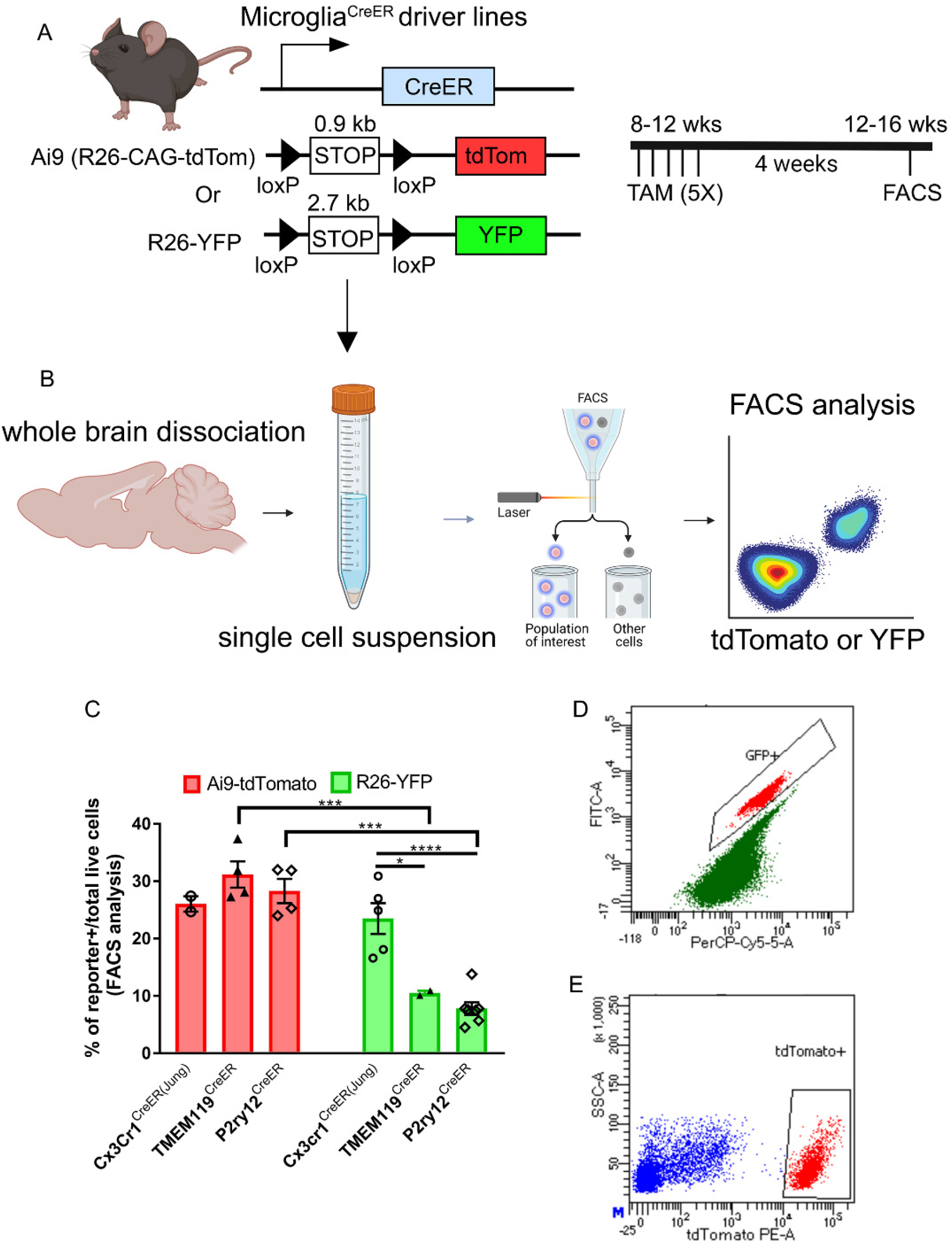
FACS analysis of reporter-positive cells in total brain single cell resuspension confirms the reporter recombination efficiency differences among the three investigated CreER lines. tdTomato+ cell percentage from all three different lines show similar high-efficiency recombination and the YFP+ cells show higher recombination efficiency in the CX3CR1CreERJung-R26-YFP mice but significantly lower recombination efficiency in the P2RY12^CreER^-R26-YFP mice and TMEM119^CreER^-R26-YFP line. Each data point represents data from one animal. **p < 0.01 and ***p < 0.001, for Two way ANOVA analysis, Tukey post-hoc pairwise analysis. Data were combined from 2-3 independent cohorts of mice for each line.

**Supplementary Table 1.** Double reporter labeling of P2ry12CreER(+/WT) after TAM Quantification of the FACS analysis of the reporter+ microglia in whole brain single cell suspension prepared from the P2RY12CreER-Ai9-R26-YFP mice after TAM treatment.

| Supplementary Table 1. Double reporter labeling of P2ry12CreER(+/-WT) after TAM |  |  |  |  |
| --- | --- | --- | --- | --- |
| Cell Condition | Mouse 1 | Mouse 2 | Mouse 3 | Average |
| % of total tdTomato+ | 77.77778 | 88.09524 | 81.69014 | 82.52105 |
| % of total YFP+ | 45.29915 | 35.71429 | 43.66197 | 41.55847 |
| % of YFP+:tdTomato- | 22.22222 | 11.90476 | 18.30986 | 17.47895 |
| % of YFP-:tdTomato+ | 54.70085 | 64.28571 | 56.33803 | 58.44153 |
| % of YFP+:tdTomato+ | 23.07692 | 23.80952 | 25.35211 | 24.07952 |
| *All percentages are calculated by dividing the number of each category of cells with the total reporter positive cells from each animal. |  |  |  |  |

